# High-fidelity reproduction of visual signals by electrical stimulation in the central primate retina

**DOI:** 10.1101/2022.05.24.493162

**Authors:** Alex R. Gogliettino, Sasidhar S. Madugula, Lauren E. Grosberg, Ramandeep S. Vilkhu, Jeff Brown, Huy Nguyen, Alexandra Kling, Paweł Hottowy, Władysław Dąbrowski, Alexander Sher, Alan M. Litke, E.J. Chichilnisky

## Abstract

Electrical stimulation of retinal ganglion cells (RGCs) with electronic implants provides rudimentary artificial vision to people blinded by retinal degeneration. However, current devices stimulate indiscriminately and therefore cannot reproduce the intricate neural code of the retina. Recent work has demonstrated more precise activation of RGCs using focal electrical stimulation with multi-electrode arrays in the peripheral macaque retina, but it is unclear how effective this can be in the central retina, which is required for high-resolution vision. This work probes the neural code and effectiveness of focal epiretinal stimulation in the central macaque retina, using large-scale electrical recording and stimulation *ex vivo*. The functional organization, light response properties, and electrical properties of the major RGC types in the central retina were mostly similar to the peripheral retina, with some notable differences in density, kinetics, linearity, spiking statistics and correlations. The major RGC types could be distinguished by their intrinsic electrical properties. Electrical stimulation targeting parasol cells revealed similar activation thresholds and reduced axon bundle activation in the central retina, but lower stimulation selectivity. Quantitative evaluation of the potential for image reconstruction from electrically-evoked parasol cell signals revealed higher overall expected image quality in the central retina. An exploration of inadvertent midget cell activation suggested that it could contribute high spatial frequency noise to the visual signal carried by parasol cells. These results support the possibility of reproducing high-acuity visual signals in the central retina with an epiretinal implant.

**Significance Statement:** Artificial restoration of vision with retinal implants is a major treatment for blindness. However, present-day implants do not provide high-resolution visual perception, in part because they do not reproduce the natural neural code of the retina. Here we demonstrate the level of visual signal reproduction that is possible with a future implant by examining how accurately responses to electrical stimulation of parasol retinal ganglion cells (RGCs) can convey visual signals. Although the precision of electrical stimulation in the central retina was diminished relative to the peripheral retina, the quality of expected visual signal reconstruction in parasol cells was greater. These findings suggest that visual signals could be restored with high fidelity in the central retina using a future retinal implant.

## Introduction

Electronic neural implants have the potential to translate our evolving scientific understanding of the nervous system into clinical restoration of sensory and motor functions. An important example is epiretinal implants, which electrically activate retinal ganglion cells (RGCs), causing them to send artificial visual signals to the brain. Although this technology is already a primary treatment for blindness resulting from severe photoreceptor degeneration (Weiland et al., 2011, 2016; Goetz and Palanker, 2016), present-day devices provide limited restoration of vision. One likely reason for this is that all currently available implants activate RGCs of different types simultaneously and indiscriminately, creating artificial neural signals that do not mimic the temporally precise and cell type specific neural code of the retina (Roska and Meister, 2013). A salient example is the simultaneous activation of ON and OFF cells at the same location, which likely sends conflicting visual information to the brain.

A potential solution would be to develop a high-resolution implant that can precisely reproduce neural activity in the diverse RGC types, which are interspersed on the surface of the retina. Previous *ex vivo* work in the peripheral primate retina has shown that epiretinal electrical stimulation with high-density multi-electrode arrays can activate individual RGCs of the major types with single-cell, single-spike resolution (Sekirnjak et al., 2008; Jepson et al., 2013, 2014b), and that the distinct cell types can be distinguished by their recorded electrical signatures (Richard et al., 2016; Madugula et al., 2022a; Zaidi et al., 2022), raising the possibility of reproducing the neural code using epiretinal stimulation. More recently, a stimulation algorithm was developed to optimize the quality of vision restoration in conditions when stimulation is focal and precisely calibrated but imperfect (Shah et al., 2019b, 2022). However, it remains unclear whether these approaches can be applied effectively to the central retina, the principal target for electronic implants, because the properties and functional organization of the diverse RGC types and the achievable specificity of electrical stimulation in areas of high RGC density are not well understood.

Here, we test how effectively electrical stimulation can reproduce the neural code of two major RGC types in the central retina, and perform a direct comparison to the peripheral retina. First, using large-scale multi-electrode recordings and visual stimulation, we probe the functional organization, light responses, spiking and electrical properties of four numerically-dominant RGC types – ON parasol, OFF parasol, ON midget, OFF midget (Dacey, 2004) – in the raphe region of the central macaque monkey retina (4.5-20° eccentricity). Next, we record and stimulate ON parasol and OFF parasol cells in the raphe and demonstrate that these two cell types can be identified and distinguished from midget cells based solely on their recorded electrical signatures, and can be activated with high spatial and temporal precision. Third, we use responses to electrical stimulation in parasol cells to estimate the visual signal reproduction that is possible with a central-targeting implant that algorithmically optimizes electrical stimulation (Shah et al., 2019b, 2022). Although the selectivity of stimulation was lower in the central compared to the peripheral retina, the unwanted activation of axons was reduced, and the overall expected image quality from parasol cells was substantially higher. These results support the possibility of high-fidelity vision restoration in the central retina with an epiretinal implant.

## Results

To understand the quality of vision restoration that could be achieved with an electrical implant in the central retina, we performed recording and stimulation *ex vivo* in the macaque monkey retina with large-scale multi-electrode arrays and determined how accurately visual stimuli can be represented in electrically-evoked RGC activity.

### Identification of the major RGC types in the central retina

To determine the functional organization of major RGC types in the central retina, light responses were obtained from regions of macaque retina 1-4.5 mm from the fovea along the temporal horizontal meridian, in the *raphe* region (Vrabec, 1966), where few multi-electrode recordings have been performed (Grosberg et al., 2017). Recordings from the more commonly studied peripheral retina (5-12 mm temporal equivalent eccentricity) were also obtained for comparison. The preparation was visually stimulated with white noise while recording spikes from hundreds of RGCs simultaneously, using an electrode array with 512 electrodes and 30 µm or 60 μm pitch (Litke et al., 2004). The spike-triggered average (STA) stimulus was computed for each cell, summarizing the spatial, temporal and chromatic properties of the light response (Chichilnisky, 2001; Chichilnisky and Kalmar, 2002). Cell types were then classified by clustering based on the spatial receptive field (RF) obtained from the STA, the time course of the STA (Fig. 1), and the spike train autocorrelation function (Field et al., 2007; Rhoades et al., 2019).

**Figure 1.**
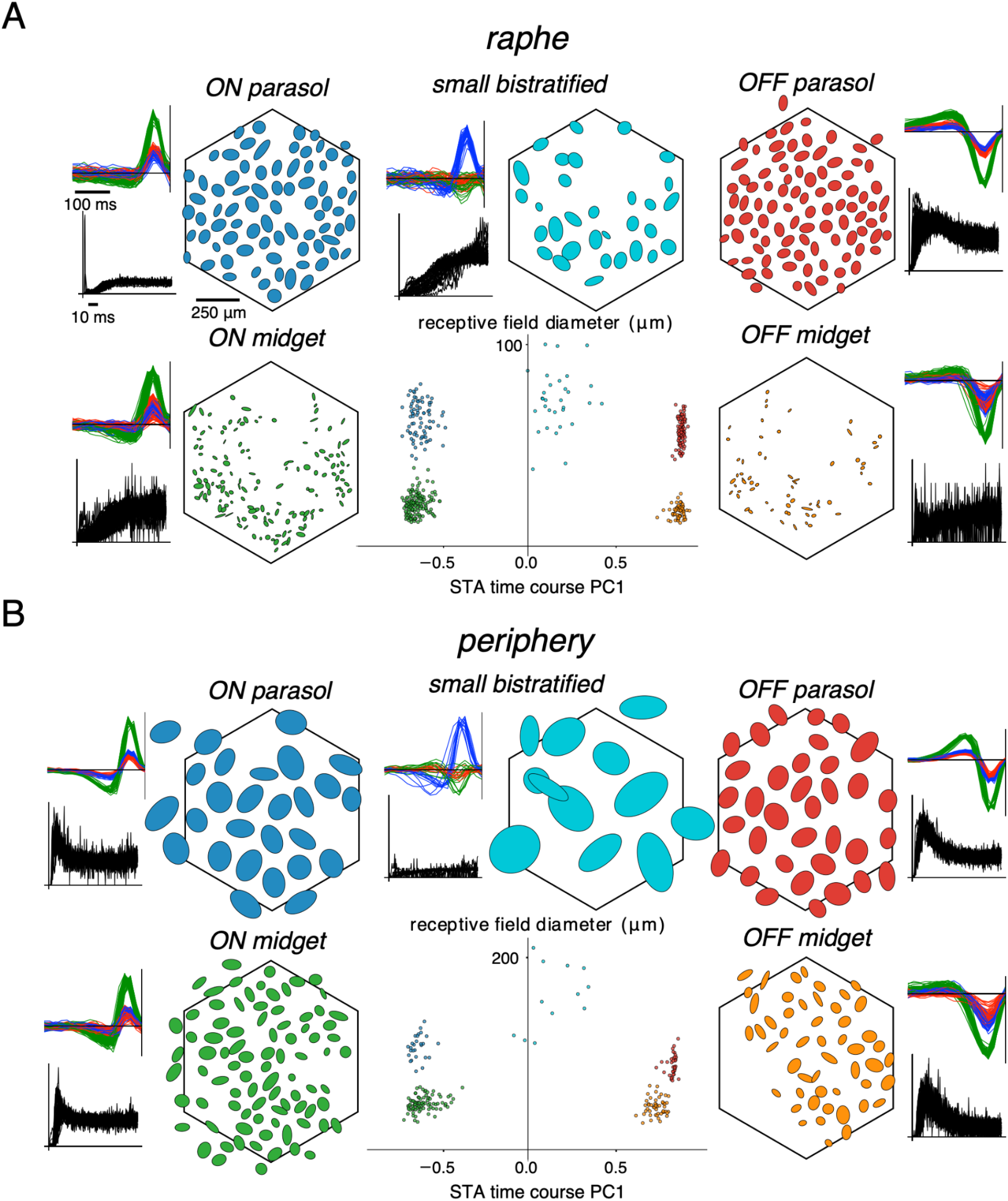
Identification of major retinal ganglion cell (RGC) types using light response and spiking properties in the raphe and peripheral retina. A. Example recording from the raphe (3.5 mm eccentricity). Clusters of receptive field (RF) diameter and the first principal component (PC1) of the spike-triggered average (STA) time course (and occasionally the PC1 of the spike train autocorrelation function, not shown) were used to separate and identify known functionally-distinct cell types. The RF mosaics, STA time courses and spike train autocorrelation functions are shown for each cell type. The ellipses denote the 2σ boundary of a 2D Gaussian fit to the spatial component of the STA. Note: other cell types were also observed, forming different clusters, but for clarity, data only for the five numerically-dominant RGC types -- ON parasol, OFF parasol, ON midget, OFF midget, and small bistratified -- are shown. Black hexagonal outline denotes the approximate location of the electrode array. B. Same as A, but for an example recording in the peripheral retina (8.5 mm temporal equivalent eccentricity).

This approach revealed the five numerically dominant primate RGC types -- ON parasol, OFF parasol, ON midget, OFF midget, and small bistratified (Fig. 1A) – which are readily identified by their light response characteristics and densities relative to other types (Dacey, 2004; Rhoades et al., 2019). Recordings from ON and OFF parasol cells were consistently nearly complete, as evidenced by the mosaic organization of their RFs (Fig. 1). Recordings of ON and OFF midget cells and small bistratified cells were less complete (Fig. 1), particularly in the raphe region (Fig. 1A). As expected, the spatial densities of these five cell types in the raphe (Fig. 1A) were higher than those of the corresponding cell types in the peripheral retina (Fig. 1B).

### Neural code of the major RGC types in the central retina

Visual signals in RGCs are often modeled using a simple cascade model, consisting of a spatiotemporal linear filter, a nonlinearity, and Poisson spiking statistics (Rodieck, 1965; Meister and Berry, 1999; Chichilnisky, 2001; Pillow et al., 2008). To compare visual processing in the raphe and periphery, this linear-nonlinear-Poisson (LNP) model was fitted to RGC responses to white noise (Chichilnisky, 2001; Chichilnisky and Kalmar, 2002), and the spatial RF, response time course and contrast-response relationship from the fitted models were examined (Fig. 2). The spatial RFs of RGCs in the raphe were on average smaller than those of their peripheral counterparts for both parasol (Fig. 2C, left, Mann-Whitney U test, *p* < 1×10^-10^ for ON and OFF) and midget cells (Fig. 2C, Mann-Whitney U test, *p* < 1×10^-10^ for ON and OFF), as expected from the smaller dendritic fields and higher density in the central retina (Watanabe and Rodieck, 1989; Dacey and Petersen, 1992). In addition, the RF overlap (see Methods) of nearest neighboring ON and OFF parasol cells was significantly higher in the raphe (Fig. 2D, Mann-Whitney U test, *p* = 0.002 for ON, *p* < 1×10^-10^ for OFF). The response time course was slower in central than peripheral parasol cells (Fig. 2E, (Sinha et al., 2017), Mann-Whitney U test, *p* < 1×10^-10^ for ON and OFF) with a more pronounced ON-OFF asymmetry (Chichilnisky and Kalmar, 2002). The time courses of central ON midget cells were significantly slower than those of peripheral ON midget cells (Fig. 2E, Mann-Whitney U test, *p* < 1×10^-10^). However, no significant difference between central and peripheral OFF midget cell time courses was observed (Mann-Whitney U test, *p* = 0.35), perhaps due to the small number of recorded OFF midget cells in the raphe. The contrast-response relationship was more linear (see Methods) in raphe parasol cells (Fig. 2F, Mann-Whitney U test, *p* = < 1×10^-10^ for ON and OFF) and midget cells (Mann-Whitney U test, *p* = 7.05×10^-10^ for ON, *p* < 1×10^-10^ for OFF), with a weaker ON-OFF asymmetry (Chichilnisky and Kalmar, 2002).

**Figure 2.**
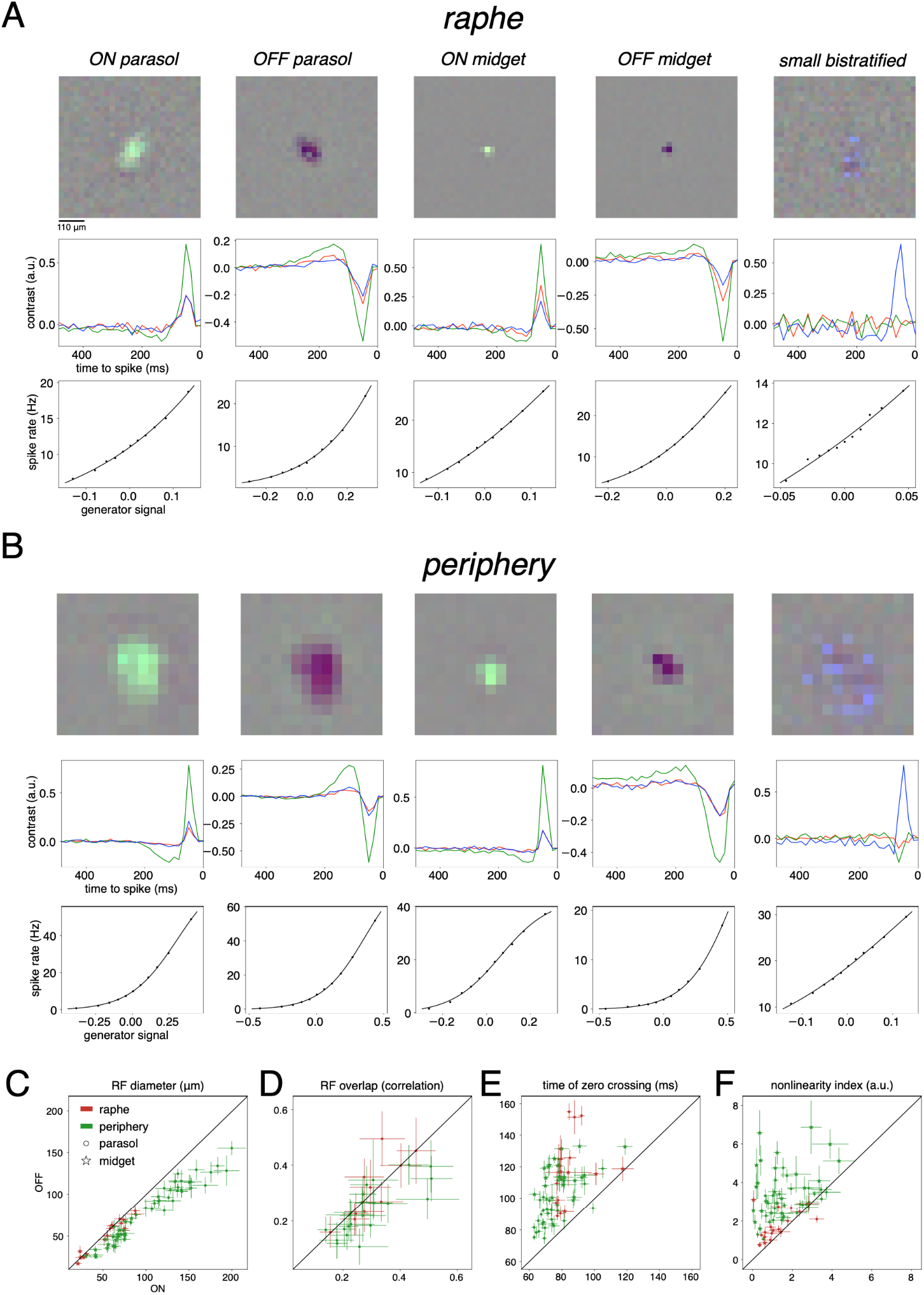
Linear-nonlinear Poisson (LNP) model comparison between raphe and peripheral RGCs. A. Example spatial component of the spike-triggered average (STA, top), STA time course (middle) and contrast-response relationship (bottom) for each of the five numerically-dominant cell types in a single raphe preparation. B. Same as A but for a peripheral preparation. C-F: Comparisons of receptive field (RF) diameter (C), RF overlap (D, see Methods), time of zero crossing of STA time course (E) and nonlinearity index extracted from the contrast-response relation (F, see Methods) between raphe and peripheral ON and OFF parasol and midget cells. Each data point represents the median value (+/- 1 median absolute deviation) from a single preparation.

To test whether the spike timing structure of central versus peripheral RGCs differs, the autocorrelation function of the spike train obtained during white noise stimulation was examined. Peripheral ON parasol cells tended to have an early peak in the autocorrelation at ∼2-5 ms, indicative of frequent doublet firing (Fig. 3A, left), compared to the more gradual increase over tens of milliseconds for raphe ON parasol cells (Fig. 3A, left). Compared to peripheral cells, the normalized early spike count (1-25 ms, see Methods) was significantly lower in raphe ON parasol cells (Fig. 3B, left, Mann-Whitney U test *p* < 1×10^-10^). Raphe and peripheral OFF parasol cell autocorrelations exhibited the same general shape, but in the raphe the autocorrelations had a significantly higher spike rate within the range of 1-100 ms (Fig. 3A, left), which was captured by the higher normalized spike count (Fig. 3A right, Mann-Whitney U test *p* < 1×10^-10^, see Methods). These findings reveal potential differences in intrinsic excitability between central and peripheral parasol cells or their synaptic inputs (see Discussion).

**Figure 3.**
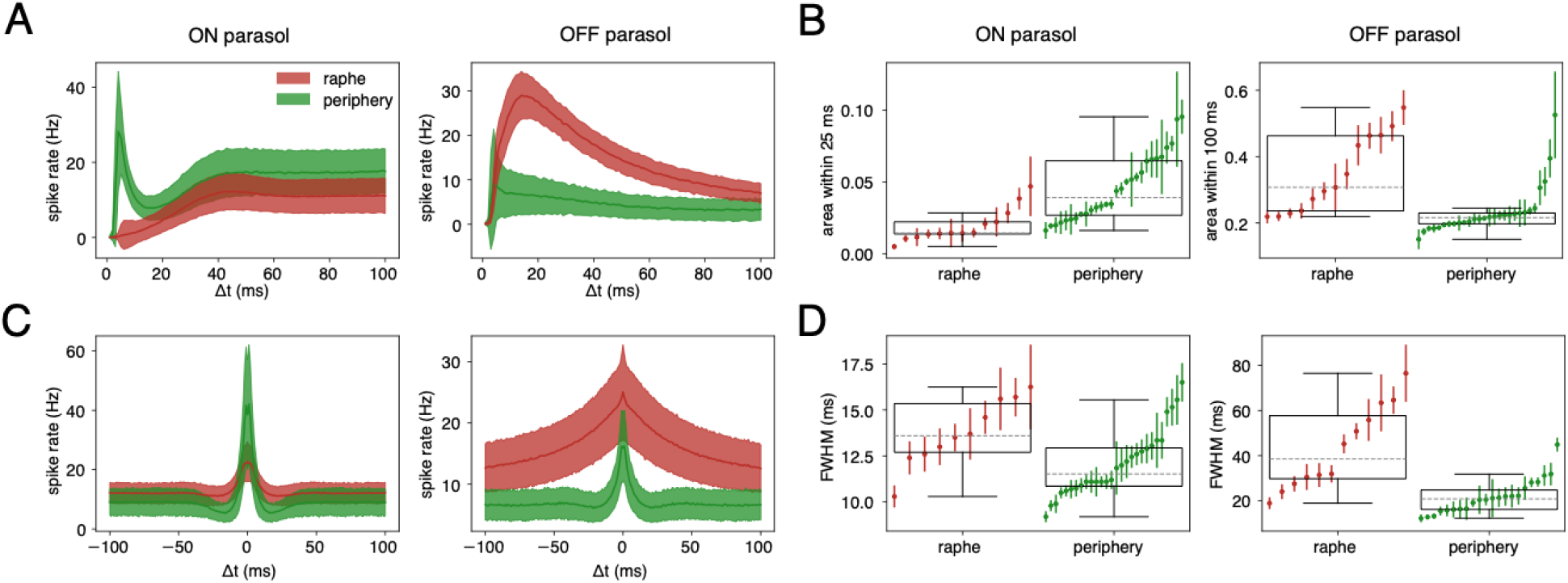
Spiking statistics and correlated firing in ON and OFF parasol cells in the raphe and the periphery. A. Mean (+/- 1 standard deviation) spike train autocorrelation functions for ON (left) and OFF (right) parasol cells in one peripheral and one raphe preparation. B. Probability of spike within 1-25 ms for ON parasol cell autocorrelations (left) and 1-100 ms for OFF parasol cell autocorrelations (see Methods). Each data point represents the median (+/- 1 median absolute deviation) value in a single preparation. C. Mean (+/- 1 standard deviation) nearest homotypic neighbor spike train cross-correlation functions for ON (left) and OFF (right) parasol cells in one peripheral and one raphe preparation. D. Full width at half maximum (FWHM, see Methods) of nearest homotypic neighbor cross-correlation functions. Each data point represents the median (+/- 1 median absolute deviation) value in a single preparation. In each box plot, the box denotes the interquartile range, the dashed gray line denotes the median value of all points in the plot, and the bottom and top whiskers denote the first quartile minus 1.5 times the interquartile range and the third quartile plus 1.5 times the interquartile range, respectively.

To test whether concerted firing between neighboring RGCs differs in the central versus peripheral retina, cross-correlation functions of spike trains obtained during white noise stimulation were computed between homotypic pairs of nearest-neighbor ON and OFF parasol cells (Mastronarde, 1983; Shlens et al., 2006, 2009; Pillow et al., 2008; Trong and Rieke, 2008). Similar to the peripheral retina, raphe ON and OFF parasol cell cross-correlations exhibited a symmetric, smoothly-decaying form peaked at the origin (Fig. 3C). However, parasol cell cross-correlations in the raphe were significantly wider (see Methods) than those in the periphery, particularly for OFF parasol cells (Fig. 3D, Mann-Whitney U test, *p* < 1×10^-10^ for ON and OFF), likely reflecting slower response time courses (Fig. 2E, see Discussion).

In summary, the neural code of major RGCs in the raphe, including light response properties (Fig. 2), spiking statistics (Fig. 3A,B), and concerted firing of neighboring RGCs (Fig. 3C,D), was largely similar to that in the peripheral retina, with some systematic differences in RF density, kinetics, nonlinearity, and firing statistics.

### Distinguishing major RGC types using intrinsic electrical features

Reproduction of the neural code with electrical stimulation requires identifying cell types in the absence of light responses (which are degraded or absent during degeneration) as well as measuring the sensitivity of each cell to electrical stimulation (Li et al., 2015; Richard et al., 2016; Shah et al., 2019a; Madugula et al., 2022a, 2022b; Zaidi et al., 2022). To probe these properties, electrical images, or the average spatiotemporal electrical footprint of a cell’s spike (Litke et al., 2004; Petrusca et al., 2007), were examined for the recorded RGCs. Electrical images of simultaneously-recorded RGCs in the raphe revealed axons that projected in one of two directions roughly 90° apart (Fig. 4A, left), consistent with the unique anatomical structure of this region (Vrabec, 1966; Grosberg et al., 2017), whereas in the periphery, the axons of RGCs all projected in the same direction toward the optic disc (Fig. 4A, right). Spike amplitudes were significantly smaller in raphe compared to peripheral parasol cells in both somatic (Mann-Whitney U test, *p* < 1×10^-10^) and axonal (Mann-Whitney U test, *p* < 1×10^-10^) cellular compartments (Fig. 4B), likely reflecting the smaller cell bodies and axon diameters in the raphe. Finally, the axon spike conduction velocity (see Methods) was significantly lower in raphe compared to peripheral parasol cells (Fig. 4C, Mann-Whitney U test, *p* < 1×10^-10^).

**Figure 4.**
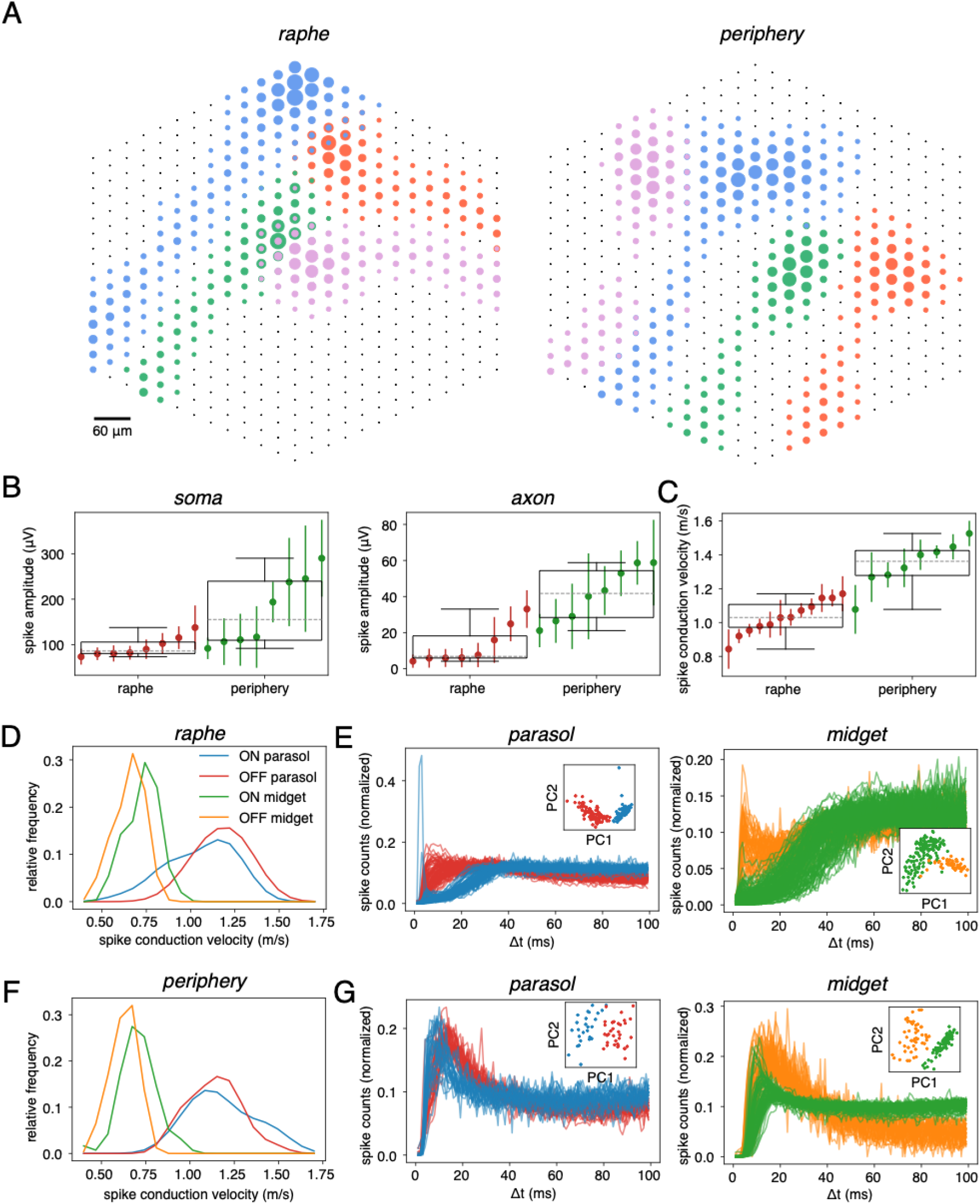
Electrical images and distinguishing cell types from intrinsic spiking and electrical features in the raphe. A. Example electrical images of four simultaneously recorded RGCs in a raphe and peripheral recording. The black points denote the locations of individual electrodes on the multi-electrode array. The collection of dots of a single color denote the electrical image of a single cell, with the size of each dot being proportional to the peak recorded voltage on that electrode during that cell’s spike. B. Maximum recorded spike amplitudes within somatic and axonal compartments within each parasol cell’s electrical image. Each data point denotes the median (+/- 1 median absolute deviation) value in a single preparation. C. Spike conduction velocities of parasol cells obtained from axonal electrodes on the electrical image (see Methods). Each data point denotes the median (+/- 1 median absolute deviation) value in a single preparation. In each box plot, the box denotes the interquartile range, the dashed gray line denotes the median value of all points in the plot, and the bottom and top whiskers denote the first quartile minus 1.5 times the interquartile range and the third quartile plus 1.5 times the interquartile range, respectively. D. Axon spike conduction velocities of ON parasol, OFF parasol, ON midget, and OFF midget cells in a single raphe preparation. For visual clarity, fitted kernel density estimates evaluated on the range of each cell type’s spike conduction velocity histogram are plotted. E. Spike train autocorrelation functions for ON and OFF parasol (left) and ON and OFF midget cells (right) in a single raphe preparation. Projections onto the first two principal components from principal components analysis (PCA) of the ON and OFF parasol (left) and ON and OFF midget cell (right) autocorrelation functions are shown as insets. Axon spike conduction velocity (D) and spike train autocorrelation functions (E) can together distinguish four of the five numerically-dominant cell types. Note that a different raphe preparation was used to show separability using the spike train autocorrelation function. F. Same as D but in the peripheral retina. G. Same as E but in the peripheral retina. A single peripheral preparation was used to show separability with axon spike conduction velocity and spike train autocorrelation function.

Raphe RGCs exhibited electrical image features and spike train statistics that can support cell type identification. Parasol cells exhibited significantly higher conduction velocities than midget cells (Fig. 4D) and could be distinguished reliably on this basis (94% classification accuracy, see Methods). Next, within the parasol and midget cell classes, principal components analysis (PCA) on the spike train autocorrelation functions (Fig. 4E) revealed two clearly-defined clusters corresponding to the ON and OFF types (Fig. 4E, insets, 100% classification accuracy for ON vs. OFF parasol cells, 94% classification accuracy for ON vs. OFF midget cells). The degree of cell type separability using these features was comparable to that in the peripheral retina (Fig. 4F,G, 97% classification accuracy for parasol-midget cells, 98% classification accuracy for ON vs. OFF parasol cells, and 100% classification accuracy for ON vs. OFF midget cells).

### Selectivity of electrical stimulation of central parasol cells

Because RGCs of different types are intermixed on the retinal surface, accurately reproducing the cell type-specific patterns of activity in the neural code (Roska and Meister, 2013) would typically require stimulation with single-cell resolution. To test whether this is possible in the raphe, spikes were identified in recorded traces immediately after application of a brief current pulse (triphasic, 0.15 ms) at each electrode individually, over a range of current levels (0.1-4.1 μA), repeated 25 times (see Methods), using the 512-electrode array with 30 μm pitch. These small currents typically directly evoke a single spike in one or more RGCs near the electrode (Sekirnjak et al., 2006, 2008; Jepson et al., 2013). For each recorded cell, the evoked spike probability was calculated at each current level, and a sigmoid fitted to response probability as a function of applied current was used to estimate the activation threshold, i.e. the current that produced a spike with probability 0.5. For each electrode, the axon bundle activation threshold was also examined, using an algorithm that identified bi-directional propagation of electrical activity to the edges of the array, a distinctive signature of axon activation ((Grosberg et al., 2017; Tandon et al., 2021), see Methods). Activation thresholds for individual parasol cells were similar in raphe and peripheral retina in both their somatic and axonal compartments (Fig. 5A). In addition, a significantly larger fraction of raphe RGCs (78%) could be activated with probability 0.5 without activating axon bundles compared to the peripheral retina (50%, resampled *p* < 1×10^-10^, Fig. 5B), consistent with a previous finding with more limited data (Grosberg et al., 2017).

**Figure 5.**
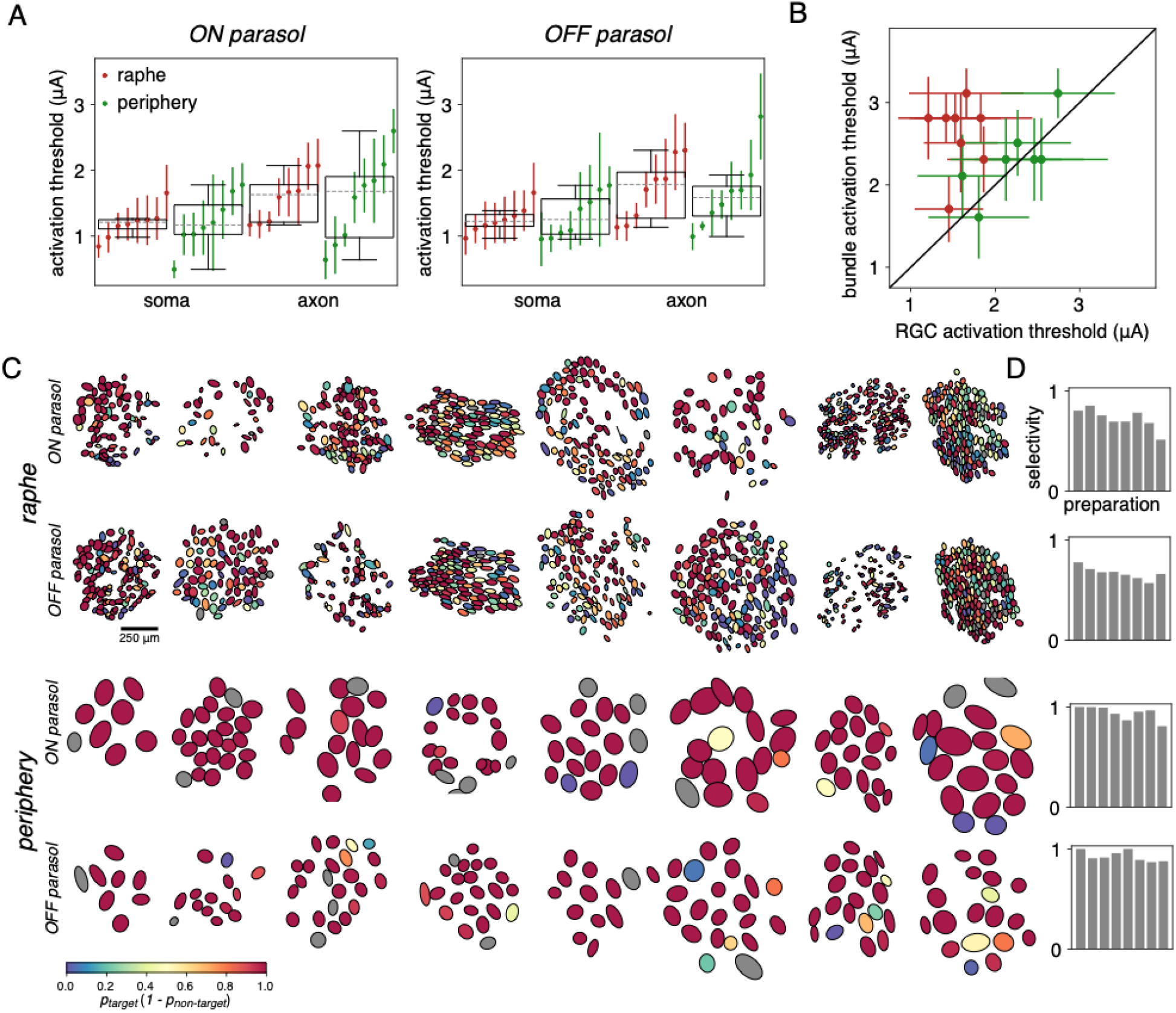
Electrical stimulation in the raphe and peripheral retina. A. Minimum somatic and axonal activation thresholds in ON parasol and OFF parasol cells. Each data point denotes the median (+/- 1 median absolute deviation) value in a single preparation. B. Minimum activation threshold versus bundle activation threshold, at each electrode. Each data point denotes the median (+/- 1 median absolute deviation) value in a single preparation. C. ON parasol cell and OFF parasol cell receptive fields (RFs) for eight raphe (top two rows) and eight peripheral retina (bottom two rows) preparations. The RF of each cell is colored according to its selectivity index (see Methods). RFs colored gray were excluded from electrical stimulation analysis because their evoked signals could not be distinguished from noise (see Methods). D. Average selectivity index values within each cell type and preparation.

To summarize how precisely individual RGCs can be activated, a selectivity index was computed for each parasol cell relative to other parasol cells: *p_target_*(*1 - p_non-target_*), where *p_target_* is the probability of firing of the target parasol cell, and *p_non-target_* is the probability of firing of the most sensitive non-target parasol cell, across stimulating electrodes and current levels (see Methods). An index of 1 indicates perfect selectivity for the target parasol cell over all non-target parasol cells that could be stimulated by the same electrode used for the target parasol cell. Across eight raphe preparations (Fig. 5C, top two rows), the average selectivity index of all ON and OFF parasol cells within the preparation ranged from 0.59-0.79 (Fig. 5D, top two panels). By comparison, the average selectivity in the peripheral retina (Fig. 5D, bottom two rows) was significantly higher (Mann-Whitney U test, *p* < 1×10^-10^ for ON and OFF parasol cells), ranging from 0.84-1.0 across preparations (Fig. 5D, bottom two panels). Greater selectivity in the periphery is expected from the lower RGC density, and is consistent with previous findings (Grosberg et al., 2017). In summary, activation of a single parasol cell without activating other recorded parasol cells is often possible in the central retina, but is less frequently achieved than in the peripheral retina.

### Inference of high-fidelity visual signal reproduction in the central retina

To determine whether electrical stimulation in the raphe can reproduce high-fidelity visual signals, the ability to reconstruct a visual image from RGC signals was evaluated (Shah et al., 2019b, 2022) in a simulation based on recorded electrically-evoked responses. First, to perform efficient image reconstruction, LNP model responses to flashed white noise images were generated, and linear reconstruction filters for each RGC were computed using least squares regression between the images and model responses (see Methods, (Warland et al., 1997; Brackbill et al., 2020)). Then, using the measured electrically-evoked responses of each cell at each stimulating electrode and each current amplitude below axon bundle activation threshold (Fig. 5), a spatiotemporal dithering strategy was used to select a pattern of electrical stimulation that would maximally decrease the expected mean-squared error (MSE) between the target image and the image reconstructed from the electrically-evoked responses (Shah et al., 2019b, 2022). The simulation produces stochastic RGC responses, and therefore, stochastic reconstructions. In this analysis, the *expected* reconstruction was examined (see Discussion).

Application of this method revealed substantial differences in reconstruction performance between the raphe and peripheral retina. Compared to the peripheral retina, the *optimal* reconstructions (i.e. the reconstructions that could in principle be achieved with perfect control over RGC activation) and the *empirical* reconstructions (i.e. the reconstructions that could be achieved, on average, with measured activation probabilities) in the raphe were both systematically higher quality than those in the peripheral retina, with sharper edges and more distinct regions that more closely matched the structure of the original image (Fig. 6A). This was evident over a range of pixel sizes (Fig. 6B). Although image reconstruction was less faithful at smaller pixel sizes in both the central and peripheral retina (Fig. 6B), image quality in the central retina was still higher at each pixel size examined.

**Figure 6.**
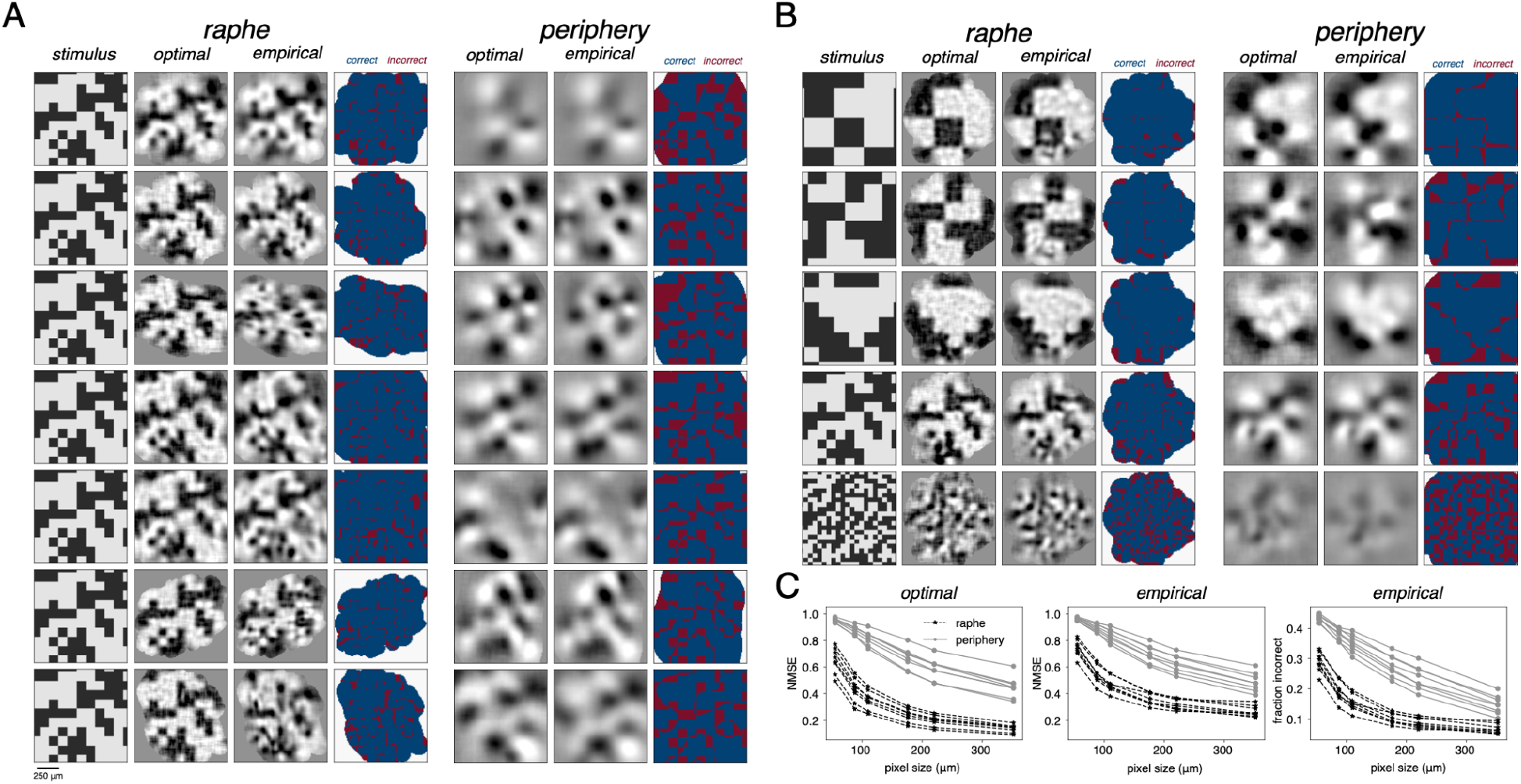
Inference of visual perception in the raphe and peripheral retina with white noise. A. Example linear reconstruction based on evoked RGC responses in individual raphe and peripheral preparations for a single white noise image (pixel size 110 μm). Each row is a distinct retinal preparation for each retinal location (two preparations per row). The first column shows the original image; the second column shows the optimal reconstruction (i.e. that achievable with perfect control over the firing of each RGC); the third column shows the reconstruction that is achievable by optimized stimulation based on recorded evoked responses ((Shah et al., 2019b, 2022), see Methods); the fourth column shows the pixels that were incorrectly reconstructed relative to the original image (red=incorrect, blue=correct). Columns 5-7 show the same as columns 2-4 but in the peripheral retina. B. Reconstruction of white noise images with different pixel sizes (352, 220, 176, 110 and 55 μm) in an example raphe and peripheral preparation. Each row is for a distinct white noise image. Columns show the same as in A. C. normalized mean-squared error (NMSE, squared error normalized by L2-norm of target image squared) in the optimal reconstructions (left) empirical reconstructions (middle) and fraction of incorrectly-reconstructed pixels in the empirical reconstructions (right). Each data point denotes the average of 15 images at each pixel size examined, for a single preparation.

The reconstructed image quality was measured by computing the normalized mean-squared error (NMSE, see Methods) between the reconstruction and original image, and the fraction of pixels with incorrect polarities in the reconstruction. In the central retina, both NMSE and the fraction of incorrect pixels were significantly lower compared to the peripheral retina, for each pixel size examined (Fig. 6C, Mann-Whitney U test, *p* < 1×10^-10^ for NMSE and fraction incorrect, pooled across all pixel sizes). However, the difference in reconstruction quality between the optimal and empirical reconstructions was significantly larger in the raphe (Mann-Whitney U test, *p* < 1×10^-10^, pooled across all pixel sizes). These findings suggest that despite the fact that stimulation is less selective in the central retina (Fig. 5C,D), it is still sufficient to produce higher quality visual signals than in the peripheral retina.

### Reconstruction of naturalistic images

The statistics of images in the natural visual world are very different from those of the white noise images tested above. To test how effectively the perception of naturalistic stimuli could be restored in the central retina using electrical stimulation, naturalistic images were reconstructed from evoked activity using a more sophisticated nonlinear reconstruction approach, as follows. Responses to flashed naturalistic images from the ImageNet database (Fei-Fei et al., 2009) were generated using the LNP model, and were used to compute optimal linear reconstruction filters. A convolutional autoencoder (CAE) was then trained to denoise the linear reconstructions generated from the filters (linear-CAE, see Methods, (Parthasarathy et al., 2017)). Using the same stimulation algorithm and procedure as above (Fig. 6, (Shah et al., 2019b, 2022)), expected reconstructions were then obtained from simulated electrically-evoked responses, based on measured single-electrode stimulation data (Fig. 5), and were enhanced using the CAE. Several aspects of the naturalistic images (distinct shapes, textures and high spatial frequency content) were more accurately captured by the raphe reconstructions than by the peripheral retina reconstructions (Fig. 7A), indicating that important features of naturalistic images can be restored in the raphe. The image quality (measured by NMSE and by SSIM, a perceptual similarity metric (Wang et al., 2004)) was significantly higher in raphe preparations (Fig. 7B, Mann-Whitney U test, *p* < 1×10^-10^ for NMSE and SSIM). In addition, similar to the white noise images (Fig. 6C), the error gap between the optimal and empirical naturalistic image reconstructions was significantly larger in raphe compared to peripheral preparations (Mann-Whitney U test, *p* < 1×10^-10^ for NMSE and SSIM), highlighting the need for further improvement to the specificity of electrical stimulation in the central retina (see Discussion).

**Figure 7.**
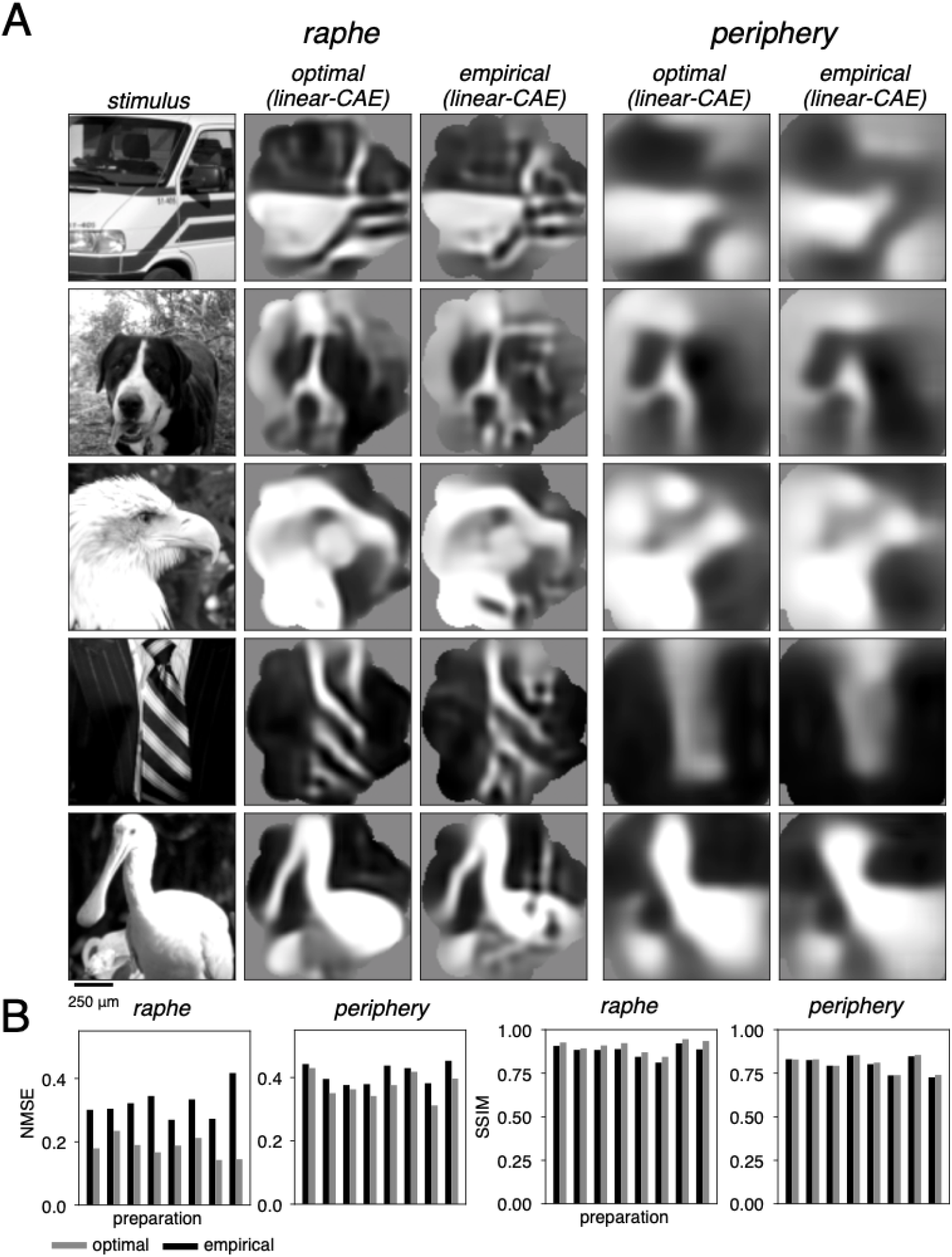
Inference of visual perception in the raphe and peripheral retina with naturalistic images. A. Reconstruction of naturalistic images from the ImageNet database (Fei-Fei et al., 2009) in an example raphe (left) and peripheral (right) preparation after image enhancement with a convolutional autoencoder (CAE, (Parthasarathy et al., 2017), see Methods). Each row corresponds to a distinct image. The first column shows the original image; the second column shows the optimal reconstruction (i.e. achievable with perfect control over the firing of each RGC); the third column shows the reconstruction that is achievable by optimized stimulation ((Shah et al., 2019b, 2022), see Methods). Columns 4 and 5 are the same as 2 and 3 but for an example peripheral preparation. Scale bar: 250 μm (1.25 degrees of visual angle). B. Optimal and empirical normalized mean-squared error (left, NMSE, squared error normalized by L2-norm of target image squared) and SSIM (right, (Wang et al., 2004)) averaged over 100 naturalistic images from the ImageNet database for each raphe and peripheral preparation. Note that in some of the peripheral preparations, the average SSIM for the empirical reconstructions is slightly higher than for the optimal reconstructions, which is possible because the CAE is trained to minimize MSE (see Methods).

### Estimation of midget cell activation

Although the majority of midget cells were not reliably identified in raphe recordings (Fig. 1A), electrical stimulation targeting parasol cells probably also activates many midget cells, because their activation thresholds are similar (Jepson et al., 2013; Madugula et al., 2022a). This inadvertent activation could introduce noise into any aspect of visual perception that is based on both parasol and midget cell signals. To estimate the extent of noise corruption, a raphe recording with partially-complete ON and OFF midget cell populations (Fig. 8A) was used to simulate complete populations of ON and OFF midget cells, including their light and electrical stimulation response properties (Fig. 8B,C, see Methods). Then, using these properties (Fig. 8B,C) and the electrical stimulus sequence targeting parasol cells (Figs. 6,7), the expected midget cell reconstruction was determined (see Methods) and summed with the expected parasol cell reconstruction to estimate the average performance of the reconstruction with added noise, for both white noise and naturalistic images. For the naturalistic image reconstructions, the same trained CAE used above (Fig. 7) was applied to denoise the reconstructions. In this simulation, the recorded responses to electrical stimulation of midget cells were not used in optimizing electrical stimulation – a worst-case analysis.

**Figure 8.**
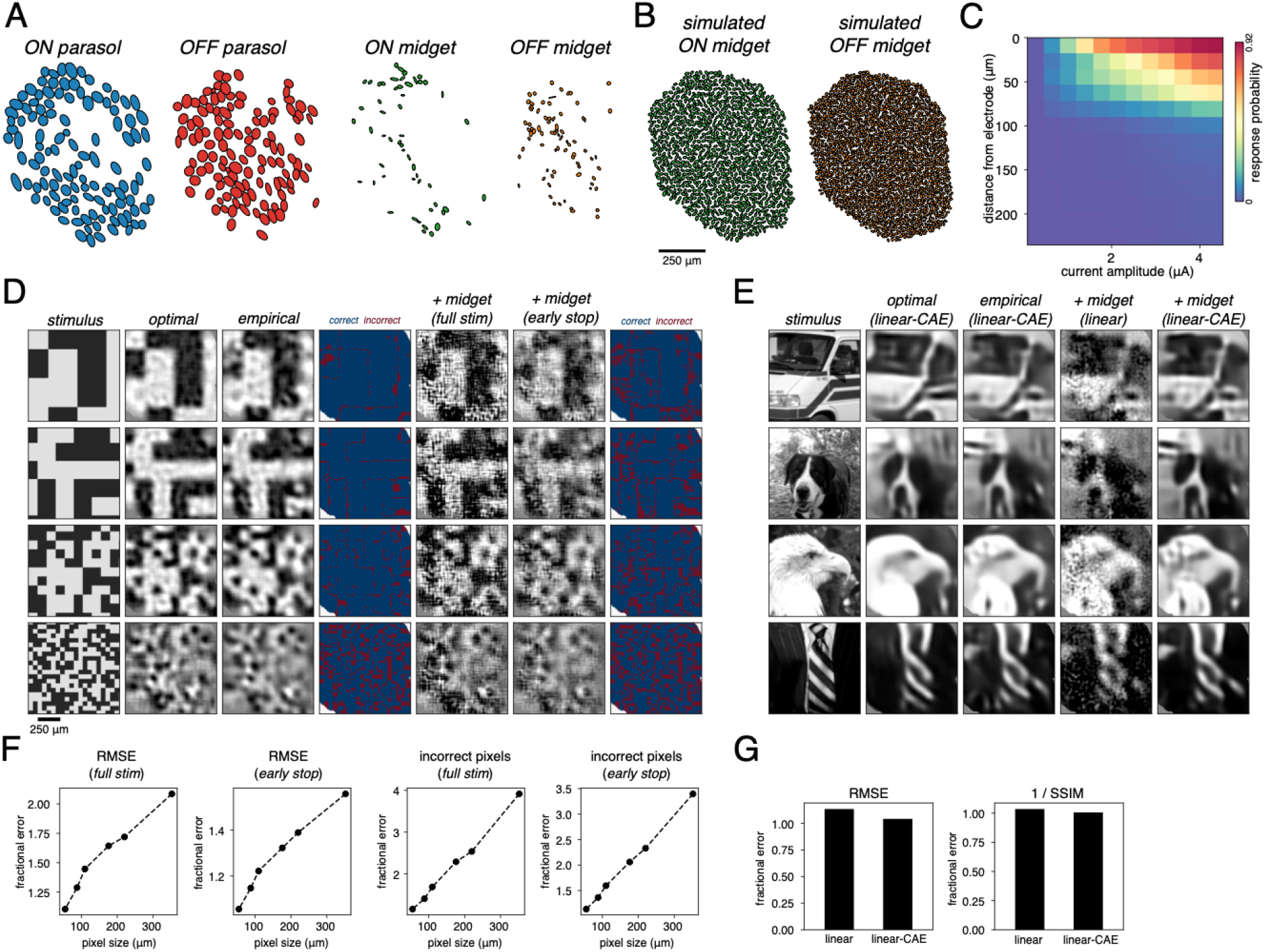
Estimation of midget cell activation. A. Receptive fields (2σ boundary from a Gaussian fit to the spatial component of the spike-triggered average, STA) of nearly complete ON parasol and OFF parasol cell populations and partial ON midget and OFF midget cell populations, in a single raphe preparation. B. Receptive fields of simulated complete populations of ON midget and OFF midget cells in the same preparation as A (see Methods). C. Response probability as a function of distance from stimulating electrode and applied current (see Methods). This relationship was determined using all of the electrical stimuli and responses from recorded midget cells in A. D. White noise reconstructions from parasol cells and the resulting reconstruction from the activation of simulated midget cells, in a single raphe preparation (A-C, pixel sizes shown: 352, 220, 110 and 55 μm). The first column shows the original image; the second column shows the optimal reconstruction (i.e. achievable with perfect control over the firing of each RGC); the third column shows the empirical reconstruction that is achievable by optimized stimulation based on recorded evoked responses ((Shah et al., 2019b, 2022), see Methods); the fourth column shows the pixels that were incorrectly reconstructed relative to the original image (red=incorrect, blue=correct); the fifth column shows the reconstruction (third column) summed with the simulated midget cell noise (see Methods); the sixth column shows the reconstruction summed with the simulated midget cell noise that can be achieved using an early stopping criterion (see Methods); the seventh column shows the pixels that were incorrectly reconstructed relative to the original image using the early stopping criterion (red=incorrect, blue=correct). E. Naturalistic image reconstructions from parasol cells and the resulting reconstruction from the activation of midget cells in the same preparation as D. The first column shows the original image; the second column shows the optimal reconstruction after application of the same trained convolutional autoencoder (CAE) as in Fig. 7; the third column shows the empirical reconstruction after application of the CAE; the fourth column shows the linear empirical parasol cell reconstruction (linear portion of column 3) summed with the linear simulated midget cell noise; the fifth column shows the resulting image after applying the CAE to the image in column 4. F. Fractional error between the white noise empirical parasol cell reconstruction (column D3) and the empirical parasol cell reconstruction summed with midget cell noise, after full stimulation (column D5) or after early stopping (column D6). Fractional error was calculated as *error_parasol+midget_/error_parasol_* where *error* is either root mean-squared error (RMSE) or fraction of incorrect pixels. Each data point denotes the average fractional error over 15 images at each pixel size. G. Fractional error between the naturalistic image empirical parasol cell reconstruction and the empirical parasol cell reconstruction summed with the midget cell noise. Fractional error was calculated using both the linear reconstructions as well as the linear reconstructions enhanced by the CAE, using both RMSE and 1 / SSIM. The bar plots denote the average fractional error over 100 naturalistic images.

Not surprisingly, inadvertent activation of midget cells added significant high spatial frequency noise to both white noise and naturalistic image reconstructions (Fig. 8D-G). For white noise image reconstructions, to reduce the magnitude of the midget cell noise, a modification of the stimulation algorithm was implemented. Normally, the algorithm chooses electrical stimuli until no further reduction in expected MSE between the parasol cell reconstruction and the target image is possible (Shah et al., 2019b, 2022). However, an early stopping criterion which takes into account an estimate of the midget cell noise (see Methods) resulted in a smaller number of electrical stimuli, strongly reducing the midget cell noise contribution (Fig. 8D,F), and thus increasing the quality of the final reconstructed image, although a significant noise corruption from midget cell activation still remained (6-56% increase in root-MSE (RMSE) and 14-240% increase in fraction incorrect pixels, Fig. 8F). For naturalistic image reconstruction, application of the CAE to the linear parasol and midget cell reconstruction removed much of the high spatial frequency noise contributed by midget cells, leaving only a small amount of noise in the final image (4% increase in RMSE and < 1% increase in 1 / SSIM, Fig. 8E,G).

The fact that the parasol signal is still recognizable in the reconstructions (Fig. 8D,E) suggests that inadvertent midget cell activation, while significant, may not be a major problem, particularly when algorithmic modifications such as early stopping are implemented or when natural image priors are taken into account, as with the CAE (see Discussion). This analysis was repeated with a second raphe recording (not shown), showing similar results: 1-13% increase in RMSE and 21-275% increase in fraction of incorrect pixels for white noise images and < 1% increase in RMSE and 1 / SSIM for naturalistic images.

## Discussion

We examined the quality of visual signals that can be restored in the raphe region of the central macaque retina with epiretinal electrical stimulation. The functional organization and light response properties of major RGC types in the raphe were similar to those of the peripheral retina (Figs. 1-3), with some notable differences, and the four numerically-dominant cell types could be distinguished using their recorded electrical features alone (Fig. 4D). Despite less precise control over electrical activation in the raphe compared to the peripheral retina (Fig. 5C,D), the unwanted activation of axons was lower in the raphe (Fig. 5B), and the estimated quality of restored visual signals in parasol cells was higher (Fig. 6A), over a range of spatial scales (Fig. 6B), and with naturalistic images (Fig. 7). These findings support the viability of targeting the raphe area for vision restoration with a high-fidelity epiretinal implant.

### Image reconstruction with parasol cell signals in the raphe

The differences in visual reconstruction from electrically-evoked signals in the raphe and periphery could not have been easily predicted on first principles alone, and thus required direct measurement. In particular, in spite of reduced stimulation selectivity (Fig. 5C,D), the fidelity of visual reconstruction was higher in the raphe (Figs. 6,7), because of the greater reconstruction resolution afforded by more cells per unit area and because of the reduced activation of axon bundles. However, the difference in performance between the optimal and empirical reconstructions was systematically larger in raphe preparations for white noise (Fig. 6) and naturalistic (Fig. 7) images, reflecting the reduced selectivity of stimulation relative to the peripheral retina (Fig. 5C,D).

Most raphe preparations exhibited higher quality reconstructions with naturalistic images compared to peripheral preparations (Fig. 7). However, one raphe preparation in particular exhibited substantially lower performance (measured by NMSE) compared to other preparations (Fig. 7B), approaching the quality of some of the peripheral preparations. This is likely due to the poor selectivity of stimulation in this preparation (Fig. 5C, rightmost raphe preparation). Multi-electrode stimulation strategies (Townshend and White, 1987; Sweeney et al., 1990; Bonham and Litvak, 2008; Martens et al., 2011) can be used to augment single-electrode stimulation, which may potentially enhance reconstruction performance. These methods have been applied previously in the peripheral retina for minimizing inadvertent neighboring RGC activation (Jepson et al., 2014a; Fan et al., 2019) as well as distant RGC activation (Nanduri, 2011) by the active avoidance of axonal stimulation (Vilkhu et al., 2021), using biophysical models of cellular activation as a theoretical basis (Rattay, 1986, 1999; Rattay and Resatz, 2004).

### Contribution of midget cell signals

Although analysis of ON and OFF parasol cells (Figs. 5,6,8) provides an indication of the visual image that can be encoded by electrical stimulation, it is not a complete representation of the visual neural code. Parasol cells constitute ∼15% of the overall RGC input to the brain (Dacey, 2004) and a lower percentage in the central retina (Marshak, 2009). Midget cells (∼50%) mediate high-acuity vision in primates, and color vision is thought to be mediated primarily by midget and small bistratified (∼5%) cells (Dacey, 2004; Field et al., 2007, 2010). Most midget and small bistratified cell spikes were not reliably identified in the raphe recordings (Fig. 1A), so their exact impact on reconstruction could not be measured. Instead, a data-driven simulation of inadvertently evoked midget cell responses to electrical stimulation when targeting the parasol cells was used to reveal a high-spatial frequency noise component that increases the error (Fig. 8D-G). However, particularly when an early-stopping criterion was implemented for white noise images (Fig. 8D,F) or when natural image priors were imposed for naturalistic images (Fig. 8E,G), much of the reconstruction from parasol cell signals was still recognizable, suggesting that unwanted midget cell activation may not be a severe problem. Nevertheless, improved spike sorting for reliably identifying spikes from all cell types, particularly the very numerous midget cells, is an important direction for future work.

The density of peripheral midget cells is comparable to that of raphe parasol cells (Fig. 1). Thus, with respect to spatial resolution, it may be that targeting central parasol cells provides a visual signal of comparable resolution to targeting peripheral midget cells. However, there are at least two other considerations. First, it may be that targeting the central retina, which a normally sighted individual uses to scan the scene with eye movements, is intrinsically important for perception. Second, midget cells could potentially subserve a different kind of perceptual experience than parasol cells. Although it is not presently clear whether peripheral midget or central parasol cells are more useful for vision restoration, future *in vivo* work in primates could be used to target the raphe parasol and peripheral midget cells separately and independently to fully understand their respective contributions to perception.

### Light response properties of raphe RGCs

Relatively little is known about differences in light response properties and intrinsic electrical properties between central and peripheral RGCs. Previous *in vivo* studies in the macaque retina have examined several chromatic, spatial and temporal response properties in parafoveal and central RGCs (Lee et al., 1989a, 1989b, 1990; Passaglia et al., 2002). In particular, differences in responses to stimuli of varying temporal frequencies were reported between central and peripheral RGCs (Lee et al., 1990; Solomon et al., 2002). A more recent *ex vivo* study of macaque RGCs (Sinha et al., 2017) directly compared differences in response kinetics between central and peripheral RGCs and reported that central cells have slower light responses, likely as a result of differences in cone photoreceptor kinetics. While the overall functional organization and light response properties appear to be similar in the central and peripheral retina (Figs. 1,2), the differences observed here build on prior work and reveal ON-OFF asymmetries – namely, slower response kinetics and a more nonlinear contrast-response in OFF versus ON cells – as well as how they compare to the asymmetries in the peripheral retina (Fig. 2E,F, (Chichilnisky and Kalmar, 2002)).

### Spiking and electrical properties of raphe RGCs

The precise origin of differences in doublet firing tendency in ON parasol cells and spike rate in OFF parasol cells between raphe and peripheral RGCs (Fig. 3A,B) is uncertain. Although it is possible that central versus peripheral parasol cells have differences in intrinsic excitability, differences in response kinetics and contrast-response (Fig. 2E,F) could produce differences in spiking statistics during visual stimulation. Future work could more thoroughly probe the origin by analyzing spontaneous activity. Differences in the width of cross-correlations (Fig. 3C,D) between nearest-neighboring central and peripheral parasol cells are likely due to response time course differences (Fig. 2E): there was a positive correlation between the width of cross-correlations and time of zero crossing in the time course across preparations (data not shown). Future work could analyze spontaneous recordings or responses to repeated white noise stimuli (Trong and Rieke, 2008; Greschner et al., 2011) to better understand circuit-level versus stimulus-driven components of correlated firing.

Spike amplitude differences between raphe and peripheral parasol cells (Fig. 4B) may reflect differences in the size of both somatic and axonal compartments (Bestel et al., 2017), which increase with eccentricity (Rodieck et al., 1985; Sanchez et al., 1986; Watanabe and Rodieck, 1989; Walsh et al., 2000; FitzGibbon and Taylor, 2012). The smaller axon diameters also may contribute to the slower axon spike conduction velocities in raphe parasol cells (Fig. 4C, (Hodgkin, 1954)), as has been demonstrated previously in the primate and cat retina (Hsiao et al., 1984; Fukuda et al., 1988).

Spiking statistics and electrical properties were sufficient to distinguish ON and OFF parasol and midget cells in the raphe (Fig. 4D,E), as has been demonstrated previously in the peripheral retina (Fig. 4F,G, (Richard et al., 2016; Madugula et al., 2022a; Zaidi et al., 2022)). However, it is unclear how effective this method will be in a degenerated retina. Previous work in rat models of photoreceptor degeneration reported altered sodium and potassium channel expression in RGCs (Chen et al., 2013), which may affect spike conduction velocity, potentially diminishing the ability to distinguish parasol from midget cells. However, a prior investigation revealed that cell type differences in spike waveforms were preserved during photoreceptor degeneration in rats (Yu et al., 2017), suggesting that electrical properties, and potentially spike conduction velocity, can still be leveraged to separate distinct cell types during degeneration. In addition, previous studies have shown differences in intrinsic RGC excitability between healthy and degenerated rat retina (Sekirnjak et al., 2011; Ren et al., 2018). Although it was reported that differences in spiking statistics between ON and OFF cells in the rat retina *increased* during degeneration (Sekirnjak et al., 2011), implying increased ability to distinguish OFF cells from ON, it is unknown whether this will apply to the primate retina or to specific cell types within the broad ON and OFF categories.

Finally, several technical limitations of this study could be addressed in future work.

1. White noise visual stimulation for light response modeling (Fig. 2) is limited because its statistics differ greatly from the statistics of naturalistic stimuli, and could elicit RGC responses that are much different than would be observed in natural viewing conditions (Heitman et al., 2016). Thus, future work could focus on performing natural scenes stimulation for response modeling in the central retina (Heitman et al., 2016; McIntosh et al., 2016), which may require models that take into account additional properties such as cell-cell interactions (Mastronarde, 1983; Pillow et al., 2008) and spatial nonlinearities (Hochstein and Shapley, 1976; Shah et al., 2020).
2. Pixel-wise MSE, although a standard and intuitive objective function to minimize loss between target and reconstructed images, does not fully capture the quality of image reconstruction relevant for vision (Wandell, 1995). Future work could optimize the perceptual similarity of reconstructed images (Shah et al., 2019b, 2022) using metrics such as SSIM (Wang et al., 2004; Parthasarathy et al., 2017; Wu et al., 2022).
3. Linear reconstruction (Warland et al., 1997; Brackbill et al., 2020), although simple and computationally tractable, is likely too simple an approximation for how the visual system extracts information from RGC spike trains to produce visual perception and behaviors. Future work could apply more sophisticated nonlinear reconstruction methods for optimizing visual perception (Parthasarathy et al., 2017; Zhang et al., 2020; Kim et al., 2021; Wu et al., 2022), such as the convolutional autoencoder used to enhance the naturalistic image reconstructions here ((Parthasarathy et al., 2017), Fig. 8A) or denoisers trained on naturalistic images (Wu et al., 2022).
4. In each reconstruction analysis, the *expected value* of the parasol cell reconstruction (Figs. 6,7) and simulated midget cell noise contribution (Fig. 8) were analyzed. A *single trial* of the parasol reconstruction resulting from the stimulation algorithm is similar to the expected value because of the variance penalty used in optimization (Shah et al., 2019b, 2022). However, a single trial of the simulated midget cell noise tends to be larger in magnitude than the expected value (not shown) because there is no regularization possible when considering responses *post hoc*. Future work could include modifications to the stimulation algorithm to minimize reconstruction error introduced by inadvertent midget cell stimulation on a single trial.
5. Several assumptions were made about how parasol and midget cell signals and natural image priors are combined to produce reconstructions. Reconstruction filters for parasol and simulated midget cells were learned separately. However, previous work has demonstrated that the cell types included when learning filters using least-squares regression strongly influence the features of any given cell’s filter (Brackbill et al., 2020). Indeed, when the parasol and simulated midget cell filters were learned together, the strength of the parasol cell filters was suppressed substantially relative to that of the simulated midget cells compared to when the filters were learned separately (not shown). Also, the CAE was trained only on linear reconstructions obtained from parasol cells (Fig. 8E,G). When the CAE was instead trained on parasol and simulated midget cell signals together, the fidelity of the denoised parasol cell reconstruction with added midget cell noise was lower (not shown). Future work could further explore how the relative weighting of signals from different cell types and image priors influences image reconstruction.

## Materials and Methods

### Experimental procedures

Eyes were obtained from terminally anesthetized macaque monkeys (*Macaca mulatta, Macaca fascicularis*) of either sex (32 males, 6 females) used by other researchers, in accordance with Institutional Animal Care and Use Committee guidelines. Immediately after enucleation, the eye was hemisected in room light and the anterior portion of the eye and the vitreous humor was removed. The posterior portion of the eye was stored in oxygenated, bicarbonate-buffered Ames’ solution (Sigma) at 31-33°C.

In infrared or dim lighting, small (∼2×2 mm) segments of retina, along with the attached retinal pigment epithelium (RPE) and choroid, were separated from the sclera. In most preparations, the retina was then separated from the RPE; in one preparation (Fig. 1B), the RPE was left attached. The retina was placed retinal ganglion cell (RGC) side down on a custom high-density multi-electrode array and a transparent dialysis membrane was used to press the retina against the array from the photoreceptor side. A total of 52 retinal preparations were used: 15 from the raphe region and 37 from the periphery. For raphe preparations, regions 1-4.5 mm (4.5-20°) eccentricity along the temporal horizontal meridian were obtained. For peripheral preparations, segments in both temporal and nasal regions of the retina were obtained, with eccentricities ranging 5-12 mm eccentricity (22-56° temporal equivalent). Once the retina was mounted on the array, it was superfused with oxygenated, bicarbonate-buffered Ames’ solution at 32-35°C.

### Visual stimulation and recording

The retina was typically stimulated with a white noise visual stimulus from a gamma-corrected cathode ray tube monitor refreshing at 120 Hz. The three monitor primaries were modulated independently for spectral variation, or in a coordinated manner for a black-and-white stimulus. The stimulus refresh interval was either 8.37, 16.74, or 33.47 ms and the size of each stimulus pixel at the photoreceptor layer ranged from ∼22-176 μm. For a single preparation (Fig. 1B), the retina was stimulated with a gamma-corrected organic light emitting diode monitor refreshing at 60 Hz. In this case, the stimulus refresh interval was 16.57 ms and the size of each stimulus pixel at the photoreceptor layer was ∼36 μm. The recordings were obtained using a custom multi-electrode stimulation and recording system with 512 electrodes (5-15 μm diameter) with either 30 μm pitch, covering an area of 0.43 mm^2^ or 60 μm pitch covering an area of 1.7 mm^2^ (Litke et al., 2004; Hottowy et al., 2008, 2012). Raw signals from the channels were amplified, filtered (43–5,000 Hz), multiplexed, digitized at 20,000 Hz and stored for off-line analysis, as described previously (Litke et al., 2004). KiloSort (Pachitariu et al., 2016) was then applied to the raw recordings to separate and identify unique cells and their spike times.

### Cell type classification

To classify distinct cell types based on their light response properties, the spike-triggered average (STA) was computed for each cell (Chichilnisky, 2001). The receptive field (RF) of each cell was estimated by determining the pixels within the STA with signal significantly above the noise level (Gauthier et al., 2009a). The temporal component of the STA was computed by summing the primaries of pixels within the RF. The spatial component of the STA was computed for each cell by taking the inner product between the time course and the full STA, and the output was fitted with a 2D Gaussian. Clusters of light response properties (RF diameter, first principal component of time course, Fig. 1) and spiking properties (first principal component of the spike train autocorrelation function, see below) were used to identify distinct cell types (Field et al., 2007). Recordings in most peripheral and some central retina preparations typically revealed the five numerically-dominant RGC types: ON parasol, OFF parasol, ON midget, OFF midget, and small bistratified. In most preparations, recordings from ON and OFF parasol cells were nearly complete, whereas recordings from ON and OFF midget and small bistratified cells were less complete (Fig. 1).

### Linear-nonlinear Poisson cascade model

To summarize the light response properties of the major RGC types, the linear-nonlinear Poisson (LNP) cascade model was fitted to responses to white noise (Chichilnisky, 2001; Chichilnisky and Kalmar, 2002). At the retinal eccentricities recorded, the RF diameters of raphe RGCs are approximately half the diameter of those in the periphery (Figs. 1,2). Therefore, to compare model parameters in the raphe and periphery, stimuli with a 22 μm pixel width were used in the raphe and stimuli with a 44 or 55 μm pixel width were used in the periphery. Both stimuli had a 16.74 ms refresh interval. Preparations recorded at different temperatures (32, 33, 35°C) were all used and there was a modest effect of temperature (i.e. lower temperatures resulted in slightly slower response time courses). However, the effects reported here were present at all temperatures (albeit with different magnitudes), and thus are not attributable to temperature differences.

First, the RF diameter of each cell was determined by fitting a 2D Gaussian to the spatial component of the STA (see above) and calculating the geometric mean of each standard deviation parameter multiplied by 2. To summarize the response time course of each cell, the STA time course of each cell was normalized (L2) and fitted with a difference of cascades of low pass filters of the form 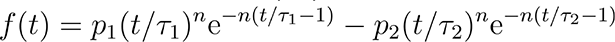 where *f(t)* is the time course, *t* is the time index, and 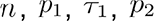 and 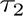 are free parameters (Chichilnisky and Kalmar, 2002). Then, the time of zero crossing of the time course was estimated by numerically solving for the root of the fitted function using the bisection method. Last, the contrast-response relationship for each cell was estimated from the STA, stimulus and spike train (Chichilnisky, 2001; Chichilnisky and Kalmar, 2002) and fitted with a smooth function of the form *f(x)* = *αL(βx+γ)*, where *x* is the generator signal, *L* is the logistic cumulative distribution function, and *α, β* and *γ* are free parameters. The *nonlinearity index* for each cell was defined as the logarithm of the ratio of *f*′(*x*_+*σ*_) and *f*′(*x*_+*σ*_) where *f*′(*x*) is the first derivative of the contrast-response relation at *x* (computed numerically using the central difference method) and *x_σ_* is the 1 standard deviation generator signal value estimated from the distribution of all generator signal values for that cell during the recording.

### Analysis of spike train autocorrelation functions

The autocorrelation function of the spike trains from white noise visual stimulation were computed for each parasol and midget cell. Briefly, spike times were binned with 1 ms precision and a histogram of pairwise spike time differences (ranging from 1 to 500 ms) was calculated and then the spike counts within each bin were then converted to a spike rate. To determine differences in firing between raphe and peripheral parasol cells, the autocorrelation was normalized to unit sum and the sum of the first 25 elements (1-25 ms) for ON parasol cells and first 100 elements (1 - 100 ms) for OFF parasol cells was calculated. These quantities provide estimates of the probability that, after the occurrence of a spike, a second spike occurs within 25 and 100 ms, respectively.

### Analysis of spike train cross-correlation functions

The cross-correlation functions of spike trains obtained during white noise visual stimulation were computed between all homotypic pairs of ON parasol and OFF parasol cells. Briefly, spike times were binned with 1 ms precision and a histogram of pairwise spike time differences ranging from −1000 to 1000 ms (0 ms inclusive) was calculated, and then the spike counts within each bin were converted to a spike rate. To determine differences in correlated firing kinetics, nearest-neighboring homotypic pairs of ON parasol and OFF parasol cells were identified and an estimate of the baseline spike rate was determined by calculating the median value of the spike rate from −1000 to -995 ms time offset and 995 to 1000 ms time offset. Next, the cross-correlation was smoothed with a Gaussian kernel (σ = 5 ms), the noise estimate was subtracted, and the resulting signal was upsampled 10x. To determine the width of the cross-correlation, the full width at half maximum (FWHM) was computed.

### Analysis of receptive field overlap

To determine the RF overlap between nearest-neighboring homotypic pairs of ON and OFF parasol cells, an estimate of each RGC’s RF was generated from its 2D Gaussian fit and the correlation coefficient was calculated between each cell’s RF and its nearest homotypic neighbor’s RF.

### Analysis of electrical images

Electrical images (EIs) were computed for each cell by averaging the voltage signal on the array during the time of each spike (Litke et al., 2004; Petrusca et al., 2007). Features of spike waveforms obtained from the EI were then used to classify them into distinct cellular compartments (Litke et al., 2004; Müller et al., 2015). In general, somatic and dendritic waveforms exhibit biphasic structure, whereas axonal waveforms exhibit triphasic structure, reflecting biophysical differences between compartments. Based on these known differences, a heuristic-based automated method (Madugula et al., 2022b) was applied to label each electrode as either “dendritic”, “somatic”, “axonal”, or “mixed” (either a superposition of somatic and dendritic or somatic and axonal waveforms, (Madugula et al., 2022b)). First, waveforms with a larger positive than negative phase were labeled as “dendritic”. Second, the waveforms at remaining electrodes were identified as “somatic”, “mixed”, or “axonal”, based on the ratio of the first positive peak to the second positive peak. Thresholds for each respective compartment were determined by two empirically observed inflection points (0.05 and 1.6 respectively) in the pooled distribution of ratios obtained from many cells and electrodes in peripheral recordings.

To determine spike amplitudes within each compartment, for each cell, electrodes recording somatic and axonal signals were determined and the maximum absolute value of the peak negative voltage on each electrode was calculated. Then, the maximum voltage over somatic and axonal electrodes was computed.

Axon spike conduction velocity was estimated from the electrical images of ON and OFF parasol and midget cells, as described previously (Madugula et al., 2022a; Zaidi et al., 2022). Briefly, axonal electrodes for each cell were identified (see above) and pairwise velocity estimates were computed by dividing the distance between each electrode by the time difference in the negative peak amplitude, for each pair of axonal electrodes. Then, a velocity estimate was computed by averaging the estimate from each pair of electrodes weighted by the product of the peak amplitudes.

### Analysis of electrical features for distinguishing cell types

Axon spike conduction velocities and the spike train autocorrelation function were used together to distinguish ON and OFF parasol and ON and OFF midget cells, to demonstrate the effectiveness of cell type classification based on intrinsic electrical features alone (Richard et al., 2016; Zaidi et al., 2022). Each preparation in this analysis (Fig. 4C and 4D,E, left) was acquired using the electrode array with 30 μm inter-electrode spacing to ensure consistent estimates of spike conduction velocities. To quantify the degree of separability between parasol and midget cell spike conduction velocities, a 1D support vector machine was fitted to the spike conduction velocity-cell type label pairs (L2-penalty with a linear kernel). The learned optimally-separating hyperplane (a point in the 1D space) was used to quantify the classification accuracy on the training data. The goal of this analysis was merely to determine the degree of separability in the data, so cross-validation with held out data was not performed.

Parasol and midget cells were highly distinguishable using spike conduction velocity (Fig. 4D,E, left). To determine if ON and OFF types within the parasol and midget cell classes could be separated by features of the autocorrelation function, the autocorrelation function was calculated (see above) and the resulting spike count vector was normalized by its L2-norm. Then, principal components analysis (PCA) was performed on a data matrix composed of either ON and OFF parasol or ON and OFF midget cell autocorrelation functions, and projections onto the first two principal components were analyzed (Fig. 4D,E middle, right, insets). To quantify the degree of separability between ON and OFF (parasol or midget) spike train autocorrelation functions, a 2D support vector machine was fitted to the projections onto the first two principal components along with the corresponding cell type labels (L2-penalty with a linear kernel). The learned optimally-separating hyperplane (a 1D line in the 2D space) was used to quantify the classification accuracy on the training data. As above, cross-validation with held-out data was not tested.

### Responses to electrical stimulation

To identify the responses to electrical stimulation of each recorded RGC, a current pulse was delivered to each electrode on the array, in a random sequence across electrodes, while recording RGC responses on all electrodes simultaneously (Jepson et al., 2013; Grosberg et al., 2017). The pulse was triphasic, positive first, and charge-balanced (50 μs per phase; relative ratios 2:-3:1). The current amplitude range tested (second phase) was 0.1-4.1 μA (∼40 amplitudes, logarithmic scale), with each amplitude repeated 25 times. In all electrical stimulation experiments, the electrode array with 30 μm inter-electrode spacing was used.

An automated spike sorting method was used to identify RGC responses to electrical stimulation. First, for each stimulating electrode, RGCs were identified as candidates for electrically evoked activity if their EIs (obtained during visual stimulation) exhibited peak amplitudes larger than 0.5 times the standard deviation of the recorded electrical noise on that electrode. Each RGC was then assigned a set of significant electrodes from its EI recording signals with power greater than two times the power of electrode noise on each electrode to be used to determine responses. Cells that did not have at least one electrode with signal power greater than 2 times the power of the electrical noise threshold were excluded from analysis because their signals could not be distinguished from noise. Then, for each current amplitude, across all RGCs and their respective significant electrodes, voltage traces recorded in the 3 ms period following stimulus application were grouped using unsupervised clustering. For each pair of clusters, signals were subtracted, aligned, and averaged to obtain differences; Typically, these difference signals reflect the firing of one or more cells in one cluster but not the other. These residuals were iteratively compared to each of the recorded spike waveforms obtained from the EIs of all RGCs and their respective significant electrodes to identify the set of RGCs contributing to the responses in each trial for each amplitude.

This algorithm produces response probabilities as a function of current level, for each RGC and stimulating electrode, which is in general monotonically increasing (Hottowy et al., 2012; Jepson et al., 2013). The relationship between current amplitude and response probability for each RGC and stimulating electrode was then fitted by a sigmoidal curve of the form 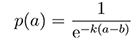, where 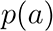 is the probability of activation, *a* is the current amplitude, and *k* and *b* are free parameters, using maximum likelihood estimation. The stimulation threshold, i.e. the current amplitude required to elicit a response probability of 0.5, was extracted from the fit (Jepson et al., 2013).

### Analysis of activation thresholds

To determine activation threshold within each compartment, for each cell, electrodes recording somatic and axonal signals were determined (see above) and the activation threshold for each electrode was extracted from the sigmoidal fit. Then, the minimum activation threshold was computed over somatic and axonal electrodes, for each cell.

### Identification of axon bundle activation thresholds

Axon bundle activation thresholds on each electrode were determined by an automated method based on a previously described algorithm (Tandon et al., 2021). The algorithm was modified to avoid bias resulting from differences in array geometries (the present work used a smaller, hexagonal array whereas the algorithm described in (Tandon et al., 2021) was developed using a larger, rectangular array) and axon spike amplitude differences between central and peripheral RGCs (Fig. 4B). For each preparation, a threshold voltage was first determined to identify electrodes that recorded significant axonal signals in response to electrical stimulation, as follows. For each RGC recorded during white noise visual stimulation, the electrodes recording axonal signals were identified as described above and the average axonal spike amplitude was determined. The median axonal spike amplitude across all recorded RGCs was computed and was taken to be the threshold voltage. Next, to determine the axon bundle activation threshold, for each stimulus current applied, electrodes were first identified as either recording a signal or not, depending on whether the recorded signal was above the threshold voltage. Activity on the array was identified as an axon bundle activation event when the collection of electrodes recording signals formed a contiguous path reaching at least two non-adjacent edges of the electrode array. The bundle activation threshold was then defined as the minimum current level at which an axon bundle activation event was evoked. Electrodes near the border of the array (outermost two rings of electrodes) were excluded from analysis because their proximity to the edge precludes the ability to unambiguously distinguish RGC activity from axon bundle activity.

### Analysis of selectivity

To summarize the selectivity of electrical stimulation, a *selectivity* metric was computed for each ON and OFF parasol cell. First, at each stimulating electrode, the response probability at each current level below axon bundle activation threshold was determined for the target cell (*p_target_*) and every other non-target parasol cells (*p_non-target_*). Then, the maximum value over current amplitudes of the quantity (*p_target_*)(1 - *p_non-target_*) was determined. This was repeated for each non-target cell, and the minimum value of (*p_target_*)(1 - *p_non-target_*) over non-target cells was determined at this stimulating electrode, to provide a worst case estimate of stimulation selectivity for this cell, at this stimulating electrode. This was repeated for each stimulating electrode, and the maximum value of (*p_target_*)(1 - *p_non-target_*) over stimulating electrodes was defined as the *selectivity index* for the cell.

### Inference of visual perception

To estimate the quality of restored vision that could be achieved with the measured responses to electrical stimulation, a recently developed stimulation algorithm (Shah et al., 2019b, 2022) was applied, in a data-driven simulation. This method assumes that the brain reconstructs visual stimuli from RGC spike trains linearly and optimally, and that responses from individual RGCs that occur within a single “integration time window” of downstream visual processing (e.g. tens of milliseconds) are summed linearly to produce perception.

In this framework, an optimal linear reconstruction filter is associated with each RGC that quantifies its contribution to the image reconstruction time it fires a spike. To compute the optimal reconstruction filter for each cell, parameters from the LNP model were used to generate responses to white noise images displayed for 100 ms. Specifically, the spike rate was simulated by taking an inner product between the STA and the stimulus and rectifying the result. The STAs were scaled such that the average parasol cell spike rate matched that observed experimentally (∼3 spikes on average per 100 ms flash in the peripheral retina). The stimuli are denoted by the matrix *S* ∈ ℝ^images x cells^ and the responses in a matrix *R* ∈ ℝ^imaes x cells^ A reconstruction matrix *W* ∈ ℝ^cells x pixels^ was computed using linear least squares regression, via the normal equations: *W* = (*R^T^R*)^−1^*R^T^S*. In this framework, the brain reconstructs an estimate of the true stimulus *S* from the RGC responses *R* by the linear operation *Ŝ* = *RW* (Warland et al., 1997). A single set of reconstruction filters was learned for a set of white noise images with varying pixel sizes (352, 220, 176, 110, 88 and 55 μm), using 10,000 training images each (60,000 images total) to test performance at a range of spatial frequencies. To produce linear reconstruction filters for naturalistic stimuli (Brackbill et al., 2020), this procedure was repeated with 60,000 grayscaled naturalistic images from the ImageNet database (Fei-Fei et al., 2009).

To determine the reconstruction that could be achieved with the responses to electrical stimulation, RGC response probabilities for each cell-electrode pair, at current levels below the threshold for axon bundle activation, were assembled in a matrix *D* ∈ ℝ^cells x (electrodes x amplitudes)^, where the columns are distinct electrical stimuli and the rows are the responses of each recorded cell. The electrical stimulus delivered at each time step was chosen such that the change in the image reconstruction due to the evoked responses of RGCs would maximally reduce the expected mean-squared error (MSE) between the reconstructed image and the target image (Shah et al., 2019b, 2022). The algorithm sequentially chooses stimuli greedily at each time step to minimize the expected MSE until no further reduction in error occurs. No two stimuli resulting in a cellular response probability of more than 0.04 were delivered within 5 ms of each other to avoid the refractory period, and each stimulation pulse was assumed to be 150 μs in duration, to be consistent with the experimental data and hardware configuration. This method was applied to a total of 90 white noise images (15 for each pixel size) and 100 naturalistic stimuli from the ImageNet database (Fei-Fei et al., 2009), all separate from the images used for training.

To approximate the optimal reconstruction, or the reconstruction that could be achieved with perfect control over the activity of each RGC, a convex solver was used to solve for the optimal nonnegative responses under squared loss between the reconstruction and the true stimulus. This response vector was then discretized by rounding each floating point response to the nearest integer value. Then, for each stimulus, the resulting response vector was applied to the reconstruction matrix to produce the approximate optimal reconstruction.

### Convolutional autoencoder

To enhance reconstructions of naturalistic images, a previously described convolutional autoencoder (CAE) (Parthasarathy et al., 2017) was applied to linear reconstructions. The CAE was developed as follows. First, linear reconstruction filters were learned for each cell using least squares regression (see above) on a set of 60,000 training images from ImageNet (Fei-Fei et al., 2009). Then, linearly reconstructed images were generated for each image in the training set using the model generated responses and reconstruction filters. This linear reconstruction was then passed through an eight layer convolutional neural network with four downsampling layers and four upsampling layers. The Adam optimizer (Kingma and Ba, 2014) with MSE loss and a learning rate of 0.0004 was used for model training. A validation set of 1,000 images from ImageNet unseen by both the linear model and the CAE was used to evaluate model accuracy. Twenty training epochs with a batch size of 32 were used, after which validation error did not substantially decrease further.

### Analysis of image reconstructions

To quantify the quality of the white noise image reconstructions, two quantities were calculated. First, the set of all relevant pixels was determined for each preparation, indicating the regions covered by the RFs of all ON and OFF parasol cells. Then, the normalized MSE (NMSE) was determined by calculating the squared error between the reconstructed and original images over this set of relevant pixels and normalizing by the squared L2-norm of the original image within this set of pixels.

Next, the fraction of reconstructed pixels with polarity opposite that of the original stimulus was calculated. A subset of the aforementioned set of relevant pixels was determined, indicating regions covered by the RFs of ON and OFF parasol cells chosen to be stimulated by the algorithm. Then, in each reconstructed image, the polarity of each pixel in the reconstruction was compared to that in the original stimulus and the fraction of pixels with opposing polarities was calculated. For the naturalistic image reconstructions, in addition to NMSE, SSIM, a perceptual similarity metric (Wang et al., 2004; Wu et al., 2022), was used to quantify image quality in a way that takes image structure into account. Note that in some of the peripheral preparations (Fig. 7B), SSIM was slightly larger for empirical naturalistic image reconstructions compared to optimal reconstructions after (but not before) applying the CAE. This is possible because the objective function for optimization in training the CAE was MSE, rather than SSIM. Separately, unlike in the raphe, the CAE had little effect on image reconstructions in some peripheral preparations because their respective linear reconstructions were coarse, due to the relatively low-density RGCs.

### Simulated midget cell receptive fields and reconstruction filters

To determine the extent of midget cell activation and the noise corruption of reconstructions that would occur when targeting only parasol cells, a raphe preparation with partial recordings of midget cells was used to simulate the electrical stimulation and visual responses of complete populations of ON and OFF midget cells (Fig. 8A,B). First, RF centers were generated from a hexagonal lattice. Ideally, the spacing of the lattice would be informed by the typical neighbor distance distribution of the midget cells. However, the midget cell populations from each raphe preparation were too sparse to reliably estimate the typical neighbor distance (Figs. 1A,7A). Since the typical neighbor distance is proportional to the typical RF diameter (Gauthier et al., 2009b), the ON (or OFF) midget cell typical neighbor distance was therefore estimated by computing the median ON (or OFF) parasol typical neighbor distance, and scaling this quantity by the ratio of median ON (or OFF) midget cell receptive field diameter to the median ON (or OFF) parasol cell receptive field diameter. This value was used to set the spacing of the lattice. The lattice locations were then jittered by adding Gaussian noise to each x and y coordinate independently, with mean 0 and standard deviation equal to 10% of the original lattice spacing. Next, using the partial populations of ON or OFF midget cells, kernel density estimates were fitted to distributions of the larger of the two (major) standard deviation parameters from the Gaussian fit to the RFs, the ratio of the major standard deviation to minor standard deviation parameters and the tilt of the receptive fields, across all ON or OFF midget cells. To avoid extreme values, the kernel density estimate was fitted to the subset of the data within the 30th and 70th percentile of each distribution. Receptive fields were then generated by randomly sampling standard deviation parameters and tilts from the fitted kernel density estimates, resulting in simulated receptive field mosaics (Fig. 8B).

Next, STAs were generated using the Gaussian parameters from the simulated receptive fields. As with the parasol cells above, the STAs were scaled such that simulated responses to white noise resulted in on average 3 spikes within the simulated ON and OFF midget cell population, consistent with empirical results from parasol and midget cells in the periphery during flashed white noise visual stimulation (data not shown). Using least squares, a set of white noise linear reconstruction filters was learned using 60,000 training images (10,000 from each pixel size) and a set of naturalistic image reconstruction filters were learned using 60,000 training images from the ImageNet database (Fei-Fei et al., 2009).

### Simulated midget cell responses to electrical stimulation

To estimate responses to electrical stimulation, recordings during single-electrode stimulation of the partial raphe midget cell populations were analyzed similarly to the parasol cells (see above) by determining response probability as a function of current amplitude and fitting a sigmoid to the resulting relationship, for each electrode-cell pair. For each cell-electrode pair, the bivariate relationship between response probability versus soma distance from electrode and current amplitude was determined. This was repeated for each cell-electrode pair and the data were pooled over all cell-electrode pairs. From the pooled data, the cell-electrode distances and current amplitudes were binned and the average response probability within each bin was calculated, providing a lookup table for a response probability, given a distance from the stimulating electrode and current amplitude (Fig. 8C).

### Estimation of midget cell noise

The electrical stimuli provided to the parasol cells for the white noise reconstruction simulation (Fig. 6A,B) were then used to generate expected responses for each simulated midget cell. Specifically, for each electrical stimulus and cell, the distance from the electrode and the current amplitude were used to determine the response probability (Fig. 8C). Over the entire stimulation sequence, the expected response for each midget cell would therefore be the sum of the probabilities from each stimulus. The resulting expected response vector was then passed as input to the linear decoder, yielding the midget cell reconstruction. Finally, this resulting reconstruction was summed with the parasol reconstruction to obtain an estimate of the expected perception when midget cells are ignored during parasol cell stimulation (Fig. 8D).

### Early stopping criterion

To reduce the magnitude of midget cell activation and consequent noise added to white noise reconstructions, an early stopping procedure was developed based on an estimate of the midget cell noise. In its original form, the stimulation algorithm used here (Shah et al., 2019b, 2022) continues to provide electrical stimuli greedily until no further reduction in expected MSE between the target stimulus and the expected parasol cell reconstruction can be achieved. In the modified procedure, the MSE between the target image and the reconstruction with the midget noise added is calculated at each electrical stimulus, and the “early stopping point” is defined as the stimulation number at which the global minimum in MSE is achieved. The exact optimal early stopping point is unknown and its estimation is directly affected by the estimate of the midget cell population, which is based on the true population, but is simulated and stochastic. To minimize potential bias in the early stopping point, 10 distinct midget cell populations were generated as above using 10 different seeds for the receptive field and electrical stimulation response generation. Then, the early stopping point for each midget cell population was determined for each of the 90 white noise stimuli and the median stopping point across the midget cell populations was calculated for each stimulus. Then, a distinct midget cell population was used for evaluation (Fig. 8B-D,F) using the early stopping points determined from the separate “training” midget cell populations.

### Experimental design and statistical analyses

In nearly all statistical tests, a two-tailed Mann-Whitney U test was performed using the statistics package in SciPy (Virtanen et al., 2020). In the statistical test comparing the fraction of RGCs that could be activated with a probability of 0.5 without activating axon bundles (Fig. 5B), a custom resampling procedure was implemented. All *p* values greater than 1×10^-10^ are reported as exact values, all others are reported as *p* < 1×10^-10^.

## Abbreviated title

Visual reconstruction in the central retina

## Acknowledgements

We thank The California National Primate Research Center, SRI International, S. Morairty, J. Carmena, J. Wallis, K. Bankiewicz, J. Horton, C. Darian-Smith, W. Newsome and T. Moore for access to primate retinas; N. Shah, E. Wu, S. Cooler, M. Zaidi, P. Vasireddy, A. Phillips, M. Breidenbach, S. Mitra and F. Rieke for useful discussions and comments on the manuscript; J. Desnoyer, R. Samarakoon, and S. Kachiguine for technical assistance; K. Mathieson for development of 30 µm pitch electrode arrays. This research was supported by NIH NIMH T32MH020016, NIH NEI F31EY033636, Fondation Bertarelli, the Stanford Neurosciences Graduate Program (ARG), NIH NEI F30-EY030776-03 (SSM), Stanford Bio-X Bowes Graduate Student Fellowship (RSV), Stanford University Vice Provost for Undergraduate Education Small Grant (HN), Polish National Science Centre grant DEC-2013/10/M/NZ4/00268 (PH), The Polish Ministry of Science and Higher Education and its grants for Scientific Research (WD), Pew Charitable Trust Scholarship in the Biomedical Sciences (AS), a donation from John Chen (AML), Stanford Medicine Discovery Innovation Award, Research to Prevent Blindness Stein Innovation Award, Wu Tsai Neurosciences Institute Big Ideas, NIH NEI R01-EY021271, NIH NEI R01-EY029247, and NIH NEI P30-EY019005 (EJC).

